# stuart: an R package for the curation of SNP genotypes from experimental crosses

**DOI:** 10.1101/2022.07.06.498978

**Authors:** Marie Bourdon, Xavier Montagutelli

**Affiliations:** Institut Pasteur, Université Paris Cité, Mouse Genetics Laboratory, Paris, France

**Keywords:** R-package, genetic analysis, SNP genotypes

## Abstract

Genetic mapping in two-generation crosses requires genotyping, usually performed with SNP markers arrays which provide high-density genetic information. However, genetic analysis on raw genotypes can lead to spurious or unreliable results due to defective SNP assays or wrong genotype interpretation. Here we introduce stuart, an open-source R package which analyzes raw genotyping data to filter SNP markers based on informativeness, Mendelian inheritance pattern and consistency with parental genotypes. Functions of this package provide a curation pipeline and formatting adequate for genetic analysis with the R/qtl package. stuart is available with detailed documentation from https://gitlab.pasteur.fr/mouselab/stuart/.

## Introduction

Genetic mapping of Mendelian or quantitative traits in inbred strains is classically achieved in two-generation crosses such as intercrosses (F2) and backcrosses (N2), in which the inheritance of the trait is compared with the genotypes at multiple genetic markers encompassing the genome map. Variations of a quantitative trait are controlled by one or more quantitative trait loci (QTL). A QTL is defined as a marker at which individuals carrying different genotypes show different average trait values. QTL mapping searches for QTLs by testing association between trait values and genotypes at markers spanning the genome map. The statistical significance of the association is expressed as logarithm of the odds (LOD) score which is calculated for each genotyped marker and, at intermediates positions, for pseudomarkers created by interval mapping, generating a LOD score curve (Broman 2001). The curve peaks at regions potentially associated with the trait. These peaks are called QTLs if they reach predefined statistical thresholds established either from general statistical models (Lander and Kruglyak 1995) or by permutation tests performed on the cross data. For each permutation, phenotypes are shuffled between individuals to break real associations, and LOD scores are calculated to identify peaks, which are all false positives. The distribution of the peak LOD scores over a large number (>1000) of permutations provides statistical thresholds: if a LOD score of 3.8 or higher is observed in 5% of the permutations, this value will be taken as the p=0.05 threshold (Doerge and Churchill 1996). QTL mapping on F2s and N2s can be conducted with R packages such as R/qtl (Broman *et al*. 2003) and R/qtl2 (Broman *et al*. 2019).

With genome sequencing, single nucleotide polymorphisms (SNPs) have become the standard across species for their very high frequency, low cost and high-throughput analysis using various genotyping platforms. In mice, several generations of Mouse Universal Genotyping Arrays (MUGA) have been developed, the most recent being GigaMUGA (143k SNPs (Morgan *et al*. 2015)) and MiniMUGA (10.8k SNPs (Sigmon *et al*. 2020)). GigaMUGA provides high-density coverage for the fine characterization of inbred strains or outbred populations such as the Diversity Outbred (Svenson *et al*. 2012), while the modest number of SNPs in MiniMUGA is largely sufficient to genotype intercrossed or backcrossed individuals. However, SNP reliability is affected by the performance of genotyping platforms and polymorphism between and within inbred strains. Spurious or unreliable mapping outputs can result from defective SNP assays or wrong genotype interpretations. Therefore, raw data obtained from genotyping services must be curated before performing genetic analyses.

Several tools exist for quality control of SNP genotyping arrays, including Illumina’s GenomeStudio. R packages such as argyle (Morgan 2016) analyze hybridization intensity signals from MUGA arrays. The simple genetic structure of two-generation crosses provides specific and efficient means for identifying spurious genotyping data, such as consistency with parental genotypes and expected Mendelian proportions. The R/qtl package includes functions to build genetic maps and check for genotype consistency (https://rqtl.org/tutorials/geneticmaps.pdf). However, this control is performed once genotypes have been imported and involves multiple steps of manual curation. To provide a more automated process of data curation before genetic analysis, we have developed stuart, an R package which implements a pipeline for automatic filtering and curation of SNP genotyping data from two-generation crosses based on simple rules. This package formats raw SNP allele calls from Illumina files into genotypes ready for importation in R/qtl. Using three intercross datasets, we illustrate the consequences of inconsistent genotypes on the estimated marker map and QTL mapping, and how the curation achieved by each function in stuart leads to trustable results.

## Materials and Methods

stuart is a tidyverse (Wickham *et al*. 2019) based R package requiring R version 3.5.0 or later. Its open source is available on Institut Pasteur’s GitLab: https://gitlab.pasteur.fr/mouselab/stuart/ and can be installed with devtools (Wickham *et al*. 2021). stuart’s vignette provides detailed descriptions of data import and of each function.

stuart imports SNP allele calls from MUGA Illumina platform or other sources using the same file format. The central object of stuart is the marker table which summarizes for each marker the alleles found in the population, the number of individuals of each genotype and the exclusion status resulting from the curation steps. stuart exports curated data to an R/qtl compatible format. The SNP annotation file used was downloaded from https://raw.githubusercontent.com/kbroman/MUGAarrays/master/UWisc/mini_uwisc_v2.csv.

Three datasets were used to test the package. This article presents the results from 176 (CC001/Unc X C57BL/6J-*Ifnar1* KO) F2 mice (dataset 1). The analysis of two other data sets, 94(C57BL/6J-*Ifnar1* KO X 129S2/SvPas-*Ifnar1* KO) F2 mice (dataset 2) and 89 (C57BL/6NCrl X CC021/Unc) F2 mice (dataset 3) is presented as supplementary data. Quantitative traits were studied in the three F2s. Phenotype distributions are presented in Supplementary Figure 1. Genotyping was performed by Neogen (Auchincruive, Scotland) with MiniMUGA on DNA prepared from tail biopsies using standard phenol-chloroform extraction. Genotype call rate was 0.927, 0.931 and 0.948 for dataset 1, dataset 2 and dataset 3, respectively. QTL mapping was performed using R/qtl. The following thresholds were used, as commonly accepted (Members of the Complex Trait Consortium 2003): p=0.05 for significant association, p=0.1 and p=0.63 for suggestive association. All figures were designed with ggplot2 (Wickham 2016) or R/qtl.

## Results and discussion

### Consequences of inconsistent genotypes

SNP data delivered by the Illumina platform are base alleles that need to be translated into genotypes for genetic analysis. From our experience on multiple two-generation crosses, we identified several types of genotype inconsistencies which were responsible for distorted marker maps and spurious QTL mapping results. Recombination fraction (RF), which measures the genetic distance between two markers, is estimated in a cross by analyzing proportion of recombinants between adjacent markers in all individuals. The map of markers calculated from the cross data should be consistent with their known positions. The R/qtl est.map() and plotMap() functions produce a graphical comparison of the two maps (Figure 1A and Supplementary Figure 2A-B). For each chromosome, the known position of each marker provided in the annotation file (left) is connected with the estimated position (right) based on observed RF. With minimally curated genotypes (exclusion of non-polymorphic markers and markers with over 50% missing genotypes), large RF were found in many instances between closely linked markers, resulting in fan-like patterns. To further describe these distortions, we computed the distribution of the ratio between the calculated and the known genetic distances between adjacent markers (Figure 1B and Supplementary Figure 2C-D; to avoid exaggerated ratios, we considered only markers with a known distance of 1cM or more). This analysis revealed two groups of markers. In dataset 1, for 43% percent of them, the ratio was below 5 and followed a Gaussian distribution with mean = 1.31 and sd = 0.77. The other markers (57%) showed a ratio between 5 and 981.87 (Figure 1B) which necessarily results from incorrect genotypes, as only a few individuals should show recombination between adjacent markers. On chromosome 1, while the known marker positions spanned ∼100 cM, the cumulated genetic distance estimated from observed RF was ∼40,000cM. As QTL mapping relies on coherent genotypes at a series of markers encompassing a genetic interval, problematic genotypes at a given marker will perturb the analysis and, in some cases, may result in peaks of the LOD score curve in the absence of true association (Cheung *et al*. 2014). Such false positives increase significance thresholds calculated by data permutation.

**Figure 1.**
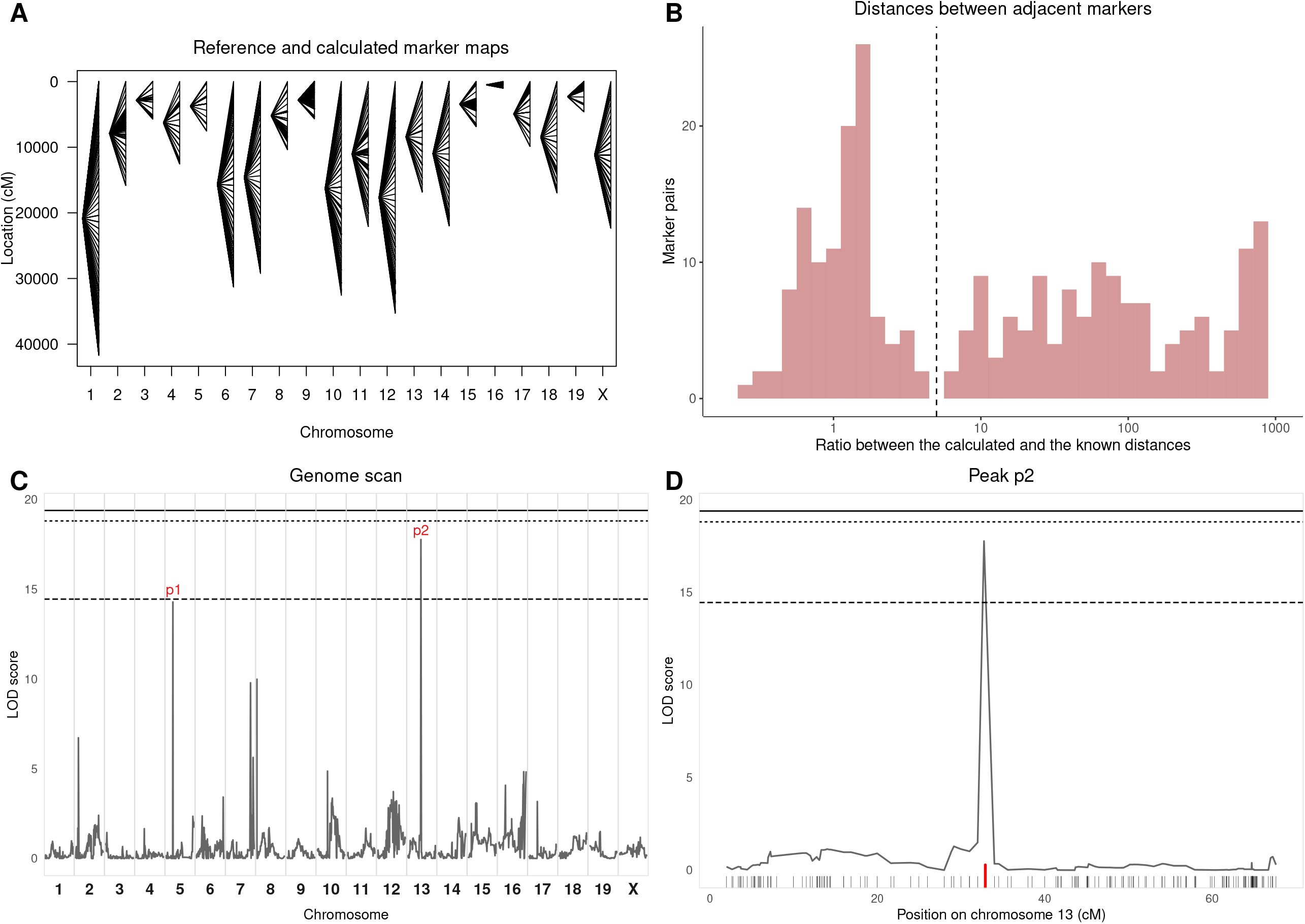
Analysis of the dataset 1 illustrating the consequences of genotyping errors and inconsistencies on QTL mapping. Non-polymorphic markers and markers with more than 50% missing genotypes were excluded to avoid excessive calculation time. A: comparison of the known marker map (left) and the genetic map estimated from observed RF (right), as calculated by est.map() and represented by plotMap() functions of R/qtl. Lines connect the positions of each marker in the two maps. The estimated map is considerably expanded because of multiple genotype inconsistencies. B: distribution of the ratio between estimated and known distances between adjacent markers. Markers with known and calculated distances below 1cM were removed as they may lead to extremely small or large ratios. The expansion of the estimated map leads to a distribution tail of high ratios. The y-axis is in logarithmic scale. 57% of markers have a ratio above 5 (dashed line). C: output of the scanone function of R/qtl showing the identification of narrow LOD score peaks. Significance thresholds are shown as plain (p=0.05), dotted (p=0.1) and dashed (p=0.63) lines. D: magnification of the scanone plot restricted to chromosome 13 (peak p2). The LOD score peak is located on one marker (red tick) distant by 1.728 cM and 1.24 cM from the proximal and distal markers, respectively, on the known marker map, but by 1001.582 cM and 1001.506 cM based on calculated RF.

These two consequences of genotyping inconsistencies are illustrated in Figure 1C and Supplementary Figure 3A-B which were obtained using the R/qtl scanone() function on a quantitative trait from the uncurated F2 datasets. For dataset 1, the p=0.05 significance threshold was estimated at 19.4 (Figure 1C), while it usually ranges between 3.3 and 4.3 depending on the inheritance model for crosses of this type and size (Lander and Kruglyak 1995). Several peaks were detected although none reached p=0.05 significance. Moreover, their narrow profile was highly unexpected in F2 crosses. Indeed, these peaks involved only one to three markers, and the LOD score curve felt abruptly between these and adjacent markers on both sides (Figure 1D), while genetic linkage between closely linked markers should result in progressive decrease of the LOD score curve on both sides of a peak (Guénet *et al*. 2015). Among the three datasets, we identified four narrow peaks reaching suggestive significance level (p<0.63): two were located at a marker with non-Mendelian allelic proportions and two were located at one to three pseudomarkers adjacent to a marker with non-Mendelian proportions (Supplementary Figure 3C-D and E-F, respectively). We identified 5 other narrow peaks (LOD score between 6.72 and 10.03) out of which four resulted from the same situations as above and one was located on a pseudomaker and a marker with non-Mendelian proportions.

Inconsistent marker maps may also originate from the wrong assignment of markers to their chromosome and position provided to the mapping program. Indeed, R/qtl developer K. Broman identified errors in MUGA arrays annotation files affecting marker positions, probe sequences mapping to several locations and unmappable markers. We recommend using K. Broman’s corrected annotation files available on GitHub. The conversion of SNP alleles (A, C, T, G) observed in second-generation individuals (SGIs) to genotypes encoded according to the parental alleles may also create genotype errors. Reference SNP alleles established for many mouse strains may be used to infer the SGI genotypes. However, we recommend genotyping individuals of the parental strains used in the cross since they could differ from the reference panel. In our example dataset, the two parental strains used in the cross showed allelic differences with their reference panel counterpart at 200 markers.

### Data control and curation performed in stuart

Although each of stuart’s functions can be called independently, we present a logical analysis workflow appropriate for two-generation crosses. Table 1 summarizes the data curation and filtering performed by each function, and the number of markers of dataset 1 retained after each step.

**Table 1.** stuart analysis pipeline and application to dataset 1.

| Steps | Function | Excluded markers | Number of markers retained |
| --- | --- | --- | --- |
| 1. Import SGI alleles from MUGA arrays | read.table()/read_tsv() | - | 11125 |
| 2. Add data from parental strain |  |  |  |
| Genotyped with SGI: make consensus | geno_strains() | - | - |
| Imported from another dataset: import and make consensus | read.table()/read_tsv(),<br>geno_strains | - | - |
| Imported from reference | read.table()/other<br>readr function<br>depending on the<br>format | - | - |
| 3. Filter on allele consistency between parents and SGI |  |  |  |
| Same set of markers between parents and SGI | mark_match() | Not present in both parents and SGI | 11125 |
| Alleles consistent between parents and SGI | mark_allele | Missing alleles in both parents<br>Not polymorphic in parents but polymorphic in SGI<br>Different alleles in parents and SGI<br>In backcrosses: homozygotes for the wrong allele<br>Optional: one parent missing or heterozygous | 10375 |
| 4. Exclude markers with high proportion of missing genotypes | mark_na() | >50% of missing genotypes by default | 9918 |
| 5. Exclude non-polymorphic markers in SGI | mark_poly() | Non polymorphic in SGI | 2738 |
| 6. Verify mendelian proportions | mark_prop() | Departure from expected Mendelian segregation (proportion of each class or statistical threshold) | 2254 |
| 7. Verify recombination fraction between markers | est.map() followed by<br>mark_estmap() | High recombination fractions with adjacent markers | 2251 |

#### Data importation

Genetic mapping requires both genotype and phenotype data. Required formats and instructions are detailed in the vignette (see example of phenotype data in Supplementary Table S1). Parental strains’ genotyping data can be loaded from the same genotyping results as the SGI, from a previous genotyping file or from a reference file. Annotation data from K. Broman can be imported directly from GitHub. The geno_strains() function formats parental genotypes from a two-allele encoding in Illumina format into a single letter encoding, and merges this data with the annotation table into a table with parental allele and marker positions.

#### Consistency between parents and SGI alleles and genotypes

Several generations of MUGA arrays have been developed (Mega, Giga, Mini), each with successive versions differing by multiple SNP markers. If parental and SGI data were produced on different versions, the marker lists must be compared to retain only common SNPs. This is achieved by the mark_match() function.

Converting alleles into genotypes requires that SGI segregate for the two parental alleles, and that each allele is found only in one parent. The aim of the mark_allele() function is to control consistency of allele’s origin at multiple levels.

First, this function excludes markers with missing data in both parents. If allele data is available for only one parent and this allele is also found in SGI, the other allele present in SGI will be assigned to the parent with missing allele. However, this imputation is not error-free since we have observed, in rare occasions, markers which alleles were identical in the parental strains but were polymorphic in the SGI (Table 2 for such SNPs in dataset 1). This situation may occur when the parental strains used in the cross have diverged from those of the reference panel, or if one parent is heterozygous. Such markers will be excluded by the mark_allele() function but they could escape detection if allele information was missing in one parent. Adding the parNH=FALSE argument to the mark_allele() function will exclude markers missing one parental allele or for which one parent is heterozygous. However, while preventing rare errors, this option will also exclude a number of truly informative markers.

**Table 2.**
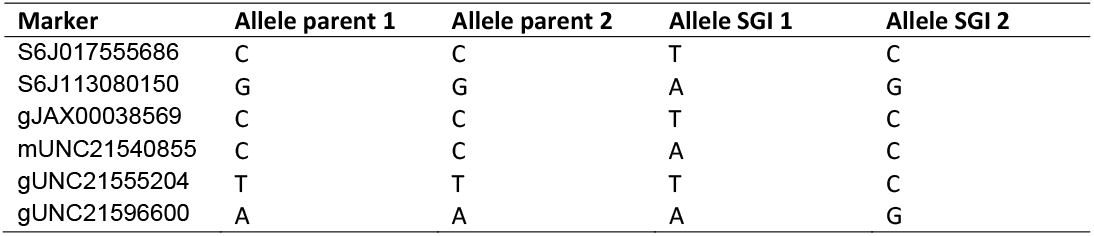
Markers of dataset 1 non polymorphic between parental strains but polymorphic in SGI.

The mark_allele() function also discards markers at which parents and SGI carry different alleles, and, for backcrosses, markers for which some SGI are homozygous for the wrong allele.

#### Non-polymorphic markers

Genetic analysis requires polymorphic markers, i.e., for which parents carry different alleles which segregate in the SGI. The mark_poly() function excludes markers for which all genotyped SGI carry the same allele, which saves computation time.

#### Missing genotypes

Reliable QTL mapping results depend on markers with medium to high rate of successful genotyping. Figure 2A shows markers distribution based on the proportion of missing genotypes. For over 95% of markers genotyping rate was above 50%. Genotyping failures may result from poor-quality genotyping assay. The mark_na() function excludes such poorly-genotyped markers.

**Figure 2.**
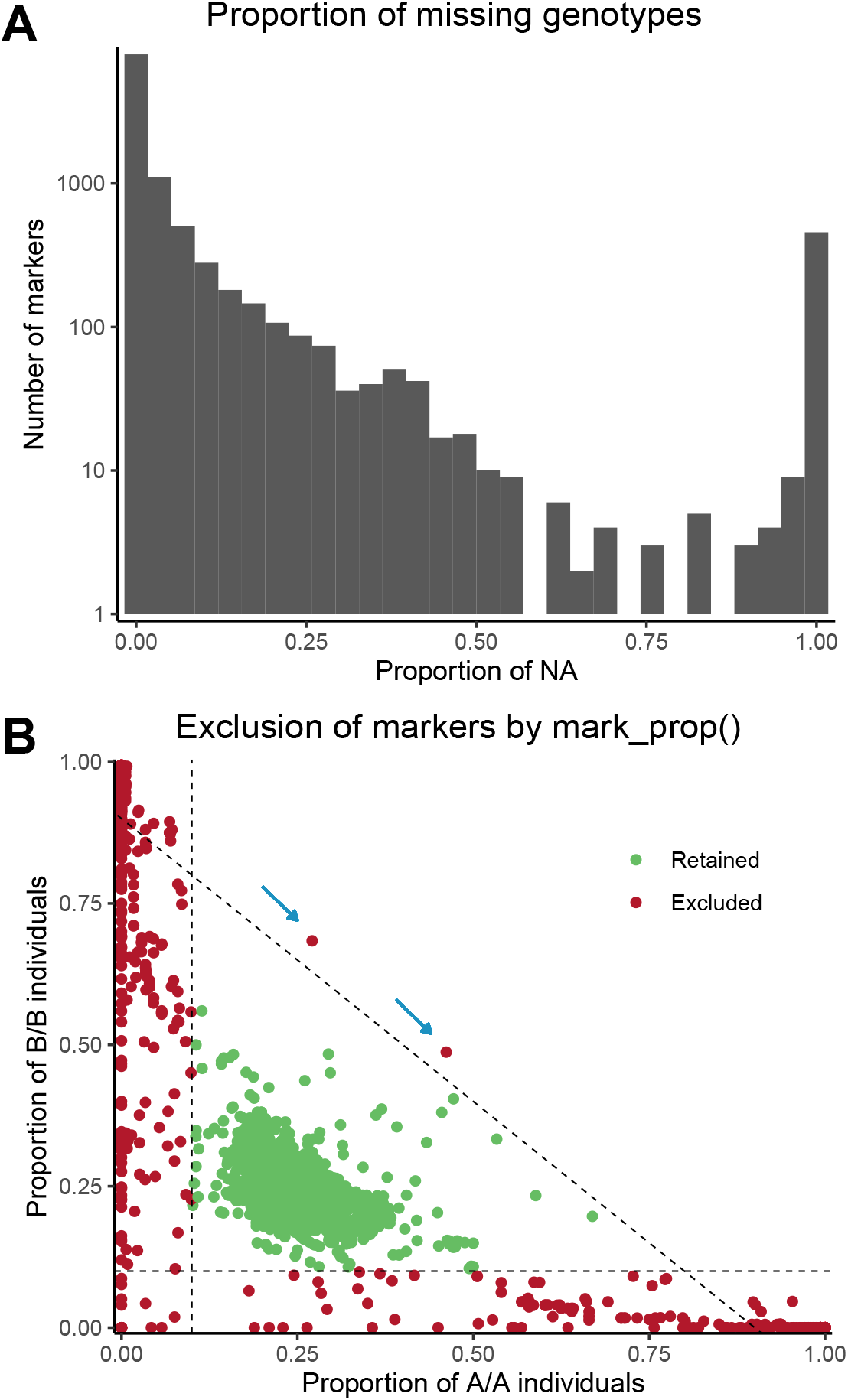
A: Distribution of the markers by their proportion of missing genotype (NA) in dataset 1. The y-axis is in logarithmic scale. 4.63% of markers have >50% missing genotypes. B: Exclusion of markers depending on genotypic proportions in dataset 1. Markers on X and Y chromosomes and mitochondrial DNA are not represented. The two axes represent the proportions of the two types of homozygous individuals in the intercross: AA and BB. Each dot represents a marker. Markers were excluded (red) if the proportion of at least one of the three genotypes (AA, AB and BB) was less than 10%, i.e. outside the triangle defined by the three dashed lines (AA=0.1, BB=0.1 and AA+BB=0.9). Blue arrows point at two markers excluded due to a proportion of heterozygotes <10%.

#### Mendelian proportions

In two-generation crosses between inbred strains, the proportions of the two or three classes of genotypes are predictable, i.e., for autosomes, 25% of each type of homozygotes and 50% of heterozygotes in an intercross, and 50% of homozygotes and 50% of heterozygotes in a backcross. Comparing the observed proportions with these expectations provides another criterion of filtering.

The mark_prop() function filters markers based either on a minimum proportion of each genotype, or on the statistically significant departure from the expected proportions (Chi2 test, with a p-value threshold). Figure 2B shows the exclusions of the autosomal markers depending on the proportion of each genotype. X chromosome genotypic proportions differ from autosomes, therefore, different arguments of mark_prop() function are used to filter X-linked markers for more precise curation.

#### Filtering report and impact on QTL mapping results

At every step, the markers filtered out are annotated in a marker table which can be exported for further inspection. The last column of Table 1 shows the number of markers retained after each step in the example dataset 1. Most of the starting markers (7180/11125 = 65%) which were eventually removed by stuart’s functions were removed by mark_poly() as non-polymorphic, a ratio expected for crosses between two standard mouse inbred strains (Frazer *et al*. 2007). mark_allele() rejected 750 markers, mark_na() 457 and mark_prop() 484. Across the three datasets, we found 1546 markers with either non-Mendelian proportions or allele inconsistencies between parental strains and SGIs. Overall, 619 of them were retained by stuart’s filtering in at least one of the other crosses, ruling out their misassignment to the genetic map. Out of the residual markers, 85 were removed from all datasets for another criterion than absence of polymorphism and were therefore considered as unreliable.

At this step, the dataset may still contain markers showing high recombination fractions with adjacent markers either for a reason not tested by the current version of stuart or due to the parameters used in mark_na() and mark_prop() functions. These markers can be identified by calculating the estimated map using R/qtl est.map()and using stuart’s mark_estmap() function which excludes markers presenting high recombination fractions with adjacent markers. Over the 3 datasets, 9 markers were removed by mark_estmap(). Five of them were retained in at least one other dataset, indicating the problem was dataset specific. Finally, for dataset 1, 2251 markers passed all steps resulting in an average genetic interval between adjacent markers lower than 2 cM, which is largely sufficient to perform QTL mapping (Darvasi *et al*. 1993). After curation, phenotype and genotype data are combined and exported in the R/qtl format using the write_rqtl() function. The qtl2convert package (Broman 2021) converts this output into the adequate format required by the more recent R/qtl2 package.

Figure 3A and Supplementary Figure 4A-B show the marker maps calculated after data curation with stuart. The known marker map and the estimated genetic map are consistent, with minimal expansions or contractions. Large ratios between the calculated and the known genetic distances between adjacent markers have been eliminated (Figure 3B, Supplementary Figure 4C-D). QTL mapping analysis on curated dataset 1 is shown on Figure 3C (to be compared with Figure 1C; see Supplementary Figure 5 for datasets 2 and 3). LOD thresholds are in the expected range for an F2, and the LOD score curve reveals broader peaks than in Figure 1B, with progressive LOD score decrease on both sides of the peak marker. One significant and three suggestive QTLs were identified on chromosomes 12 (p-value = 0.037, Figure 3D), 5 (p-value = 0.460), 10 (p-value = 0.157) and 15 (p-value = 0.244) which were not visible using non-curated data due to very high LOD score thresholds.

Being very simple to use and efficient at curating genotyping errors, stuart will facilitate the use of genotyping arrays for genetic mapping purposes in two-generation crosses, bridging the gap between raw allele data produced by SNP platforms and genetic analysis software. Moreover, its functions can be used independently to analyze inbred strains genotypes. For example, geno_strain() creates a genotype consensus between two or more individuals of the same strain suitable for further inspection, which can be useful when genotyping or regenotyping a strain of interest. Comparing genotyping results of an inbred strain after several generations of breeding with mark_allele() will readily identify variants that have emerged or been selected over time. Likewise, this function will help identifying genetic variants between substrains.

**Figure 3.**
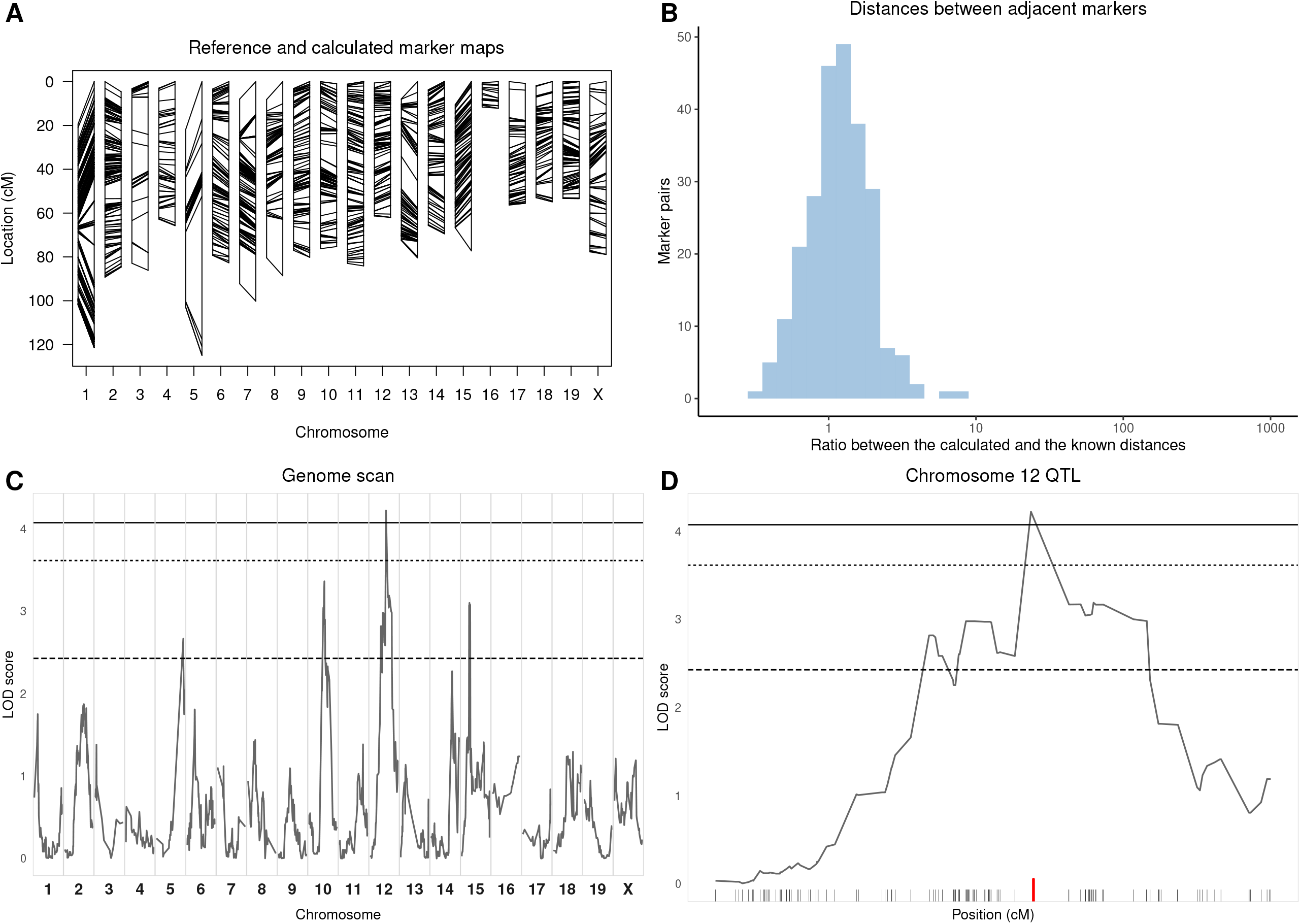
Analysis of dataset 1 after curation of genotyping data by stuart using the mark_match(), mark_allele(), mark_na(), mark_poly(), mark_prop(), and mark_estmap() functions. Refer to Figure 1 for comparison with original data. A: the estimated marker map is now consistent with the known marker map. Despite some contraction or expansion of specific intervals, the genome length of the observed marker map for each chromosome is consistent with the known map (ratio between the calculated and the known length of the genome:1.12). B: distribution of the ratio between estimated and known distance between adjacent markers. Markers with known and calculated distances below 1cM were removed as they may lead to extremely small or large ratios. Ratios are normally distributed with mean=1.33 and sd=0.81 showing consistency between the known and estimated maps. C: The LOD score curve shows several peaks, one of which is significant at P<0.05 (plain line). Note that the significance thresholds are much lower than in Figure 1C. None of the peaks shown in Figure1C were confirmed after data curation. Conversely, none of the peaks above P=0.63 (dashed line) found after data curation had been detected in Figure1C. D: magnification of the QTL peak identified on chromosome 12, showing progressive decrease of the LOD score curve over a large genetic interval. The marker with the highest LOD score is identified with a red tick.

### Web resources

The source code of the stuart package and the code used for the figures of this article are publicly available from https://gitlab.pasteur.fr/mouselab/stuart/.

## Data availability statement

All datasets used as examples in this article are available from https://gitlab.pasteur.fr/mouselab/stuart/. Dataset 1 is included in the package and can be loaded once the package is loaded (see the vignette for details). The two other datasets are available from GitLab in the “article” directory in separate folders (i.e. “data2” and “data3”). Each folder contains the genotypes of the SGIs in file “geno_dataX.csv”, the phenotypes of the SGIs in file “pheno_dataX.csv”, the parental strains’ genotypes in file “parents_dataX.csv” and the reference genotypes for the parental strains in file “ref_geno_dataX.csv”. Analysis of each cross is in each folder in an R markdown file (“dataX.Rmd”).

## Acknowledgements

We thank Elise Jacquemet of the Pasteur Institute Bioinformatics and Biostatistics HUB for helping with the use of GitLab.

## Supplementary data

## Supplementary Table 1

Format of the phenotype data. First column: individual’s number. Second column: individual’s sex. Following columns: traits and covariates (individual’s age, phenotype)

**Supplementary Figure 1**

Distribution of the quantitative phenotypes analyzed in the three datasets.

**Supplementary Figure 2**

Analysis of datasets 2 and 3 illustrating the expansion of the estimated genetic maps. Non-polymorphic markers and markers with more than 50% missing genotypes were excluded to avoid excessive calculation time. A, B: comparison of the known marker map (left) and the genetic map estimated from observed RF (right), as calculated by est.map() and represented by plotMap() functions of R/qtl in dataset 2 (A) and dataset 3 (B). Lines connect the positions of each marker in the two maps. The estimated map is considerably expanded because of multiple genotype inconsistencies. C, D: distribution of the ratio between estimated and known distance between adjacent markers. Markers with known and calculated distances below 1cM were removed as they may lead to extremely small or large ratios. The expansion of the estimated map leads to a distribution tail of high ratios. The y-axis is in logarithmic scale. 51% of the markers in the dataset 2 (C) and 16% of the markers in the dataset 3 (D) have a ratio above 5 (dashed line).

**Supplementary Figure 3**

Analysis of datasets 2 and 3 illustrating the identification of narrow LOD-score peaks. A, B: output of the scanone() function of R/qtl in the datasets 2 (A) and 3 (B) showing the identification of two narrow suggestive peaks. C: peak p1 from dataset 1 (see Figure 1C) is located on a single marker (mUNC050096588, red tick) with non-Mendelian proportions. Peak p2 from dataset 1 shows the same pattern. D: genotypes at mUNC050096588. HM1 and HM2: homozygotes; HT: heterozygotes; NA: missing genotypes. E: peak p3 from dataset 2 is located on a pseudomarker adjacent to a marker with non-Mendelian proportions (SNT111392585, red tick). Peak p4 from dataset 3 shows the same pattern. F: genotypes at SNT111392585.

**Supplementary Figure 4**

Analysis of the estimated genetic map in datasets 2 (A) and 3 (B) after curation of genotyping data by stuart. Refer to Supplementary Figure 2 for comparison with original data. The estimated marker maps are now consistent with the known marker maps with similar genome length despite local contractions and expansions (the ratio between the calculated and the known length of the genome is 1.00 for dataset 2 and 0.96 for dataset 3). The ratios between estimated and known distance between adjacent markers in dataset 2 (C) and dataset 3 (D) are now normally distributed with a mean=1.27 and a sd=0.77 for dataset 2 and a mean=1.24 and a sd=0.61 for dataset 3. The x-axis is in logarithmic scale.

**Supplementary Figure 5**

Analysis of the LOD score curve after curation of genotyping data by stuart in datasets 2 (A) and 3 (B). Refer to Supplementary Figure 3 for comparison with original data. Significance thresholds are much lower than before curation. One peak in dataset 2 is significant at P<0.05 (plain line) and none of the peaks observed before data curation (Supplementary Figure 3) were confirmed after curation with stuart. Dotted line: P=0.1. Dashed line: P=0.63.

